# Ancient antagonism between CELF and RBFOX families tunes mRNA splicing outcomes

**DOI:** 10.1101/099853

**Authors:** Matthew R Gazzara, Michael J. Mallory, Renat Roytenberg, John Lindberg, Anupama Jha, Kristen W. Lynch, Yoseph Barash

## Abstract

Over 95% of human multi-exon genes undergo alternative splicing, a process important in normal development and often dysregulated in disease. We sought to analyze the global splicing regulatory network of CELF2 in human T cells, a well-studied splicing regulator critical to T cell development and function. By integrating high-throughput sequencing data for binding and splicing quantification with sequence features and probabilistic splicing code models, we find evidence of splicing antagonism between CELF2 and the RBFOX family of splicing factors. We validate this functional antagonism through knockdown and overexpression experiments in human cells and find CELF2 represses RBFOX2 mRNA and protein levels. Because both families of proteins have been implicated in the development and maintenance of neuronal, muscle, and heart tissues, we analyzed publicly available data in these systems. Our analysis suggests global, antagonistic co-regulation of splicing by the CELF and RBFOX proteins in mouse muscle and heart in several physiologically relevant targets including proteins involved in calcium signaling and members of the MEF2 family of transcription factors. Importantly, a number of these co-regulated events are aberrantly spliced in mouse models and human patients with diseases that affect these tissues including heart failure, diabetes, or myotonic dystrophy. Finally, analysis of exons regulated by ancient CELF family homologs in chicken, and *Drosophila* suggests this antagonism is conserved through evolution.

## INTRODUCTION

Alternative splicing of pre-mRNAs is required for proper protein expression and cellular function across all metazoans (Nilsen and Graveley 2010; Kalsotra and Cooper 2011). Alternative splicing is typically controlled by binding of proteins along a nascent transcript (Fu and Ares 2014). These RNA binding proteins in turn regulate the interaction of the splicing machinery with substrate resulting in inclusion or skipping of specific exons in the final mRNA. Importantly, pre-mRNA transcripts are often bound by multiple proteins, that cooperate or compete in their regulation of the splicing machinery. Thus, the final splicing pattern of a gene in any given cell is determined by the combinatorial activity of the repertoire of RNA binding proteins associated with its pre-mRNA (Chen and Manley 2009; Fu and Ares 2014). It follows that understanding the relative expression and activity of RNA binding proteins is essential to determining how gene expression is controlled.

Two RNA binding protein (RBP) that have been particularly linked to alternative splicing in many of developmental and differentiation processes are CELF2 and RBFOX2. CELF2 is one of the six members of the human family of CUGBP, ELAV-Like Family (CELF) proteins, while RBFOX2 is a member of the RBFOX family of RBPs, which has three paralogues in humans. Both families of RBPs are strongly conserved throughout metazoa and have been well-documented to regulate tissue- and developmental-specific splicing from C. elegans to humans. In humans, the different members of the CELF and RBFOX families exhibit distinct tissue distribution and their expression is often highly regulated in a developmental-dependent manner (Dasgupta and Ladd 2011; Conboy 2016). For example, CELF2 is highly expressed in the fetal heart but show a marked reduction in expression during postnatal development (Kalsotra et al. 2008; Kalsotra et al. 2010). Aberrant re-expression of CELF2 in adult heart leads to widespread mis-regulation of splicing and aberrant cardiac function (Ladd et al. 2005; Kalsotra et al. 2008; Kalsotra et al. 2010). Overexpression of CELF2 in muscle and brain has also been show to contribute to mis-regulated splicing and pathology in several muscular dystrophies (Charlet et al. 2002; Bland et al. 2010; Dasgupta and Ladd 2011; Wang et al. 2015), while increased expression of CELF2 is normal in developing thymocytes and activated T cells and regulates splicing to promote T cell receptor expression and signaling (Mallory et al. 2011; Martinez et al. 2015). Similarly, proper expression of RBFOX2 in brain and muscle of several species is required for appropriate splicing and tissue function (Kuroyanagi et al. 2007; Gallagher et al. 2011; Singh et al. 2014). Interestingly, despite the overlap in expression and function of CELF2 and RBFOX2 in many tissues and across species, physical or functional interplay between these proteins has not been well studied.

Recently we have shown that CELF2 expression increases during thymic development and upon activation of mature T cells (Mallory et al. 2011; Mallory et al. 2015). We have also identified CELF2 dependent alternative splicing of ~100 genes in immature, mature and cultured T cells and have mapped the transcriptome-wide association of CELF2 with pre-mRNA in cultured Jurkat T cells (Mallory et al. 2015; Martinez et al. 2015; Ajith et al. 2016). These studies revealed that CELF2 functions generally in a position-dependent manner, in which binding of CELF2 upstream of a variable exon results in exon repression, while binding of CELF2 downstream of a variable exon induces its inclusion in the final mRNA (Ajith et al. 2016). Position-dependent activity has been noted for many RNA binding proteins, including RBFOX2 (Yeo et al. 2009; Fu and Ares 2014; Shankarling et al. 2014).

Interestingly, our initial analysis of CELF2 binding compared to function also indicated that many of the ~100 genes for which we observed differential splicing in CELF2 depleted cells lacked observable binding of CELF2 around the regulated exon (Ajith et al. 2016). These data suggested an additional, indirect, activity of CELF2 on splicing. Here we expand our identification of CELF2-dependent splicing in Jurkat cells, and demonstrate that over half of the exons that exhibit increased inclusion upon depletion of CELF2 lack any evidence of CELF2 binding in their vicinity. Strikingly, analysis of common features shared among CELF2-repressed exons revealed an enrichment of RBFOX binding sites in the downstream intron. This enrichment of RBFOX2 binding was observed regardless of the presence or absence of CELF2 binding. Using, RT-PCR we confirm that exons with downstream RBFOX2 binding sites are sensitive to RBFOX2 depletion in Jurkat cells. Moreover, we find that CELF2 regulates the expression of RBFOX2 in Jurkat cells at the level of mRNA stability, such that expression of RBFOX2 protein is markedly increased upon CELF2 depletion. Together this data suggests that CELF2 antagonizes RBFOX2 expression to indirectly regulate splicing, while RBFOX2 antagonizes CELF2 function when both are bound to an RNA. Importantly, by analyzing publicly available data we find evidence for antagonism between CELF and RBFOX family members across other mammalian tissues, including heart and muscle, and in chicken and flies, suggesting that antagonism between these critical splicing factors is a highly conserved mechanism for regulating splicing during development.

## RESULTS

### CELF2 regulates an extensive program of splicing in T cells both directly and indirectly

In previous studies, we used a targeted RNA-seq approach to identify ~100 cassette exons that exhibit altered splicing upon knock-down of CELF2 expression in Jurkat T cells (Mallory et al. 2015; Martinez et al. 2015; Ajith et al. 2016). As this pilot study interrogated only ~5000 exons, we sought to gain a more complete view of the consequence of CELF2 expression on the transcriptome in T cells using RNA-seq of total polyA-selected RNA from Jurkat cells. We have previously shown that CELF2 expression increases upon stimulation of Jurkat cells with the phorbol ester PMA (Mallory et al. 2015; Martinez et al. 2015). Therefore, we carried out RNA-Seq of mRNA from wildtype and CELF2-depleted Jurkat cells grown under either unstimulated or PMA-stimulated conditions. Depletion of CELF2 was done by doxycycline-inducible expression of a shRNA targeting CELFs, as we have described previously (Mallory et al. 2015; Martinez et al. 2015; Ajith et al. 2016).

Using the splicing-quantification algorithm MAJIQ (Vaquero-Garcia et al. 2016), we identified ~901 significant changes in splicing (local splice variations or LSVs) induced upon CELF2 knockdown in unstimulated cells, and ~1700 significant CELF2-dependent LSVs in stimulated cells (Fig. 1A). The larger number of CELF2-dependent LSVs in stimulated cells is consistent with the increased expression of CELF2 in these cells compared to unstimulated conditions. Despite the difference in number of CELF2-dependent LSVs in these two cell conditions, there is a significant overlap in the identity of CELF2-regulated LSVs in unstimulated and stimulated cells (Fig. 1A). Moreover, the extent of CELF2 regulation of individual LSVs in these two conditions is also highly similar (Fig. 1B), as we have observed previously (Ajith et al. 2016). Therefore, we conclude that CELF2 has a widespread and consistent impact on alternative splicing in both unstimulated and stimulated T cells. For subsequent analysis, we have thus merged the high-confidence LSVs from unstimulated and stimulated cells (i.e. those with a probability of >95% that the difference in inclusion (ΔPSI) is greater than 20%) to identify sets of cassette exons for which inclusion is increased (CELF2-repressed) or decreased (CELF2-enhanced) upon depletion of CELF2, in addition to exons that were unresponsive to CELF2 depletion (CELF2-unresponsive). This resulted in 265 CELF2-repressed, 371 CELF2-enhanced and 1,668 CELF2-unresponsive exons (Supplemental Table S1, see Methods).

**Figure 1:**
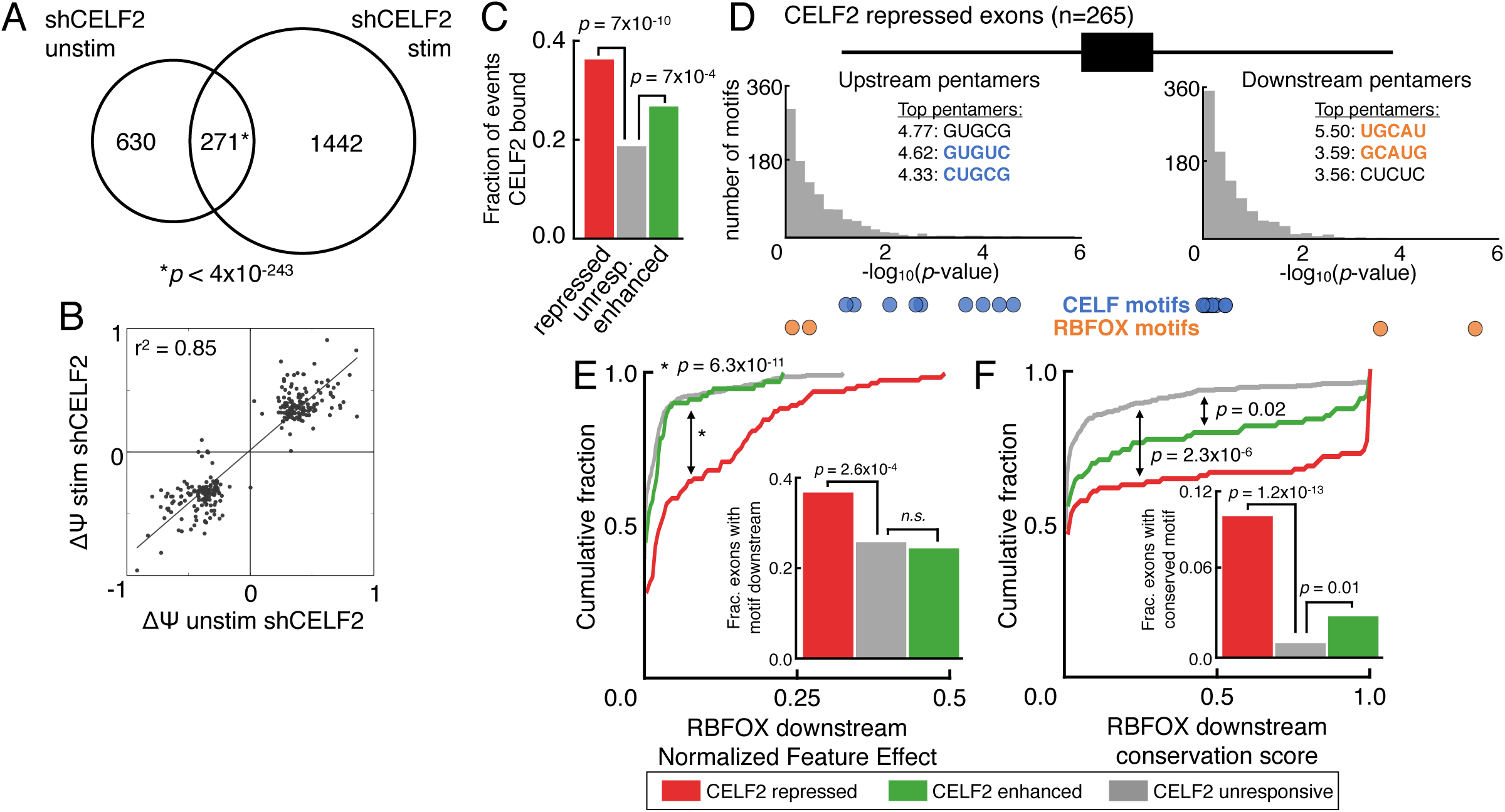
RNA-seq reveals global CELF2 regulation in T cells and sequence analysis suggests co-regulation by the RBFOX family. (A) Venn diagram of the overlap of unique, high-confidence LSVs (Vaquero-Garcia et al. 2016) that showed significantly altered splicing (IE(ΔΨ)|>20%, probability>95%) upon depletion of CELF2 by shRNA in unstimulated (left) or stimulated (right) JSL1 Jurkat T cells (Fisher's exact (FE) test, two-tailed *p* < 7×10^−192^) (B) Scatterplot comparing E(ΔΨ) values for the most changing junction from unique, co-regulated LSVs upon CELF2 depletion in unstimulated (X) and stimulated (Y) Jurkat T cells. (C) Fraction of high-confidence cassette exons containing splice site proximal CELF2 CLIP-seq peaks that were repressed by CELF2 (red, n=265), enhanced by CELF2 (green, n=373), or were unresponsive (grey, n=1668) (FE test, two-tailed) (D) Distribution of enrichment significance (-log_10_(*p*-value), hypergeometric test comparing regulated to unresponsive sets) for the occurrence of all pentamers upstream (left) or downstream (right) of CELF2 repressed exons. Top three pentamers for each are shown in the inset and motifs known to be bound by the CELF family or RBFOX family are highlighted in blue or orange, respectively. See Supplemental Table S2 for all values. (E) Cumulative distribution function (CDF) of the AVISPA predicted impact on splicing (Normalized Feature Effect, NFE; (Barash et al. 2013)) for downstream [U]GCAUG motifs for the subsets of events that contained it (Kolmogorov-Smirnov two-sample test). Fraction of each set with a downstream [U]GCAUG is given in inset with colors corresponding to the key (FE test, twotailed). (F) CDF showing the average conservation (phastCons 46way, placental mammals) for the core GCAUG motif for the subsets of events that contained it downstream of the alternative exon (KS two-sample test). Fraction of each set with a perfectly conserved downstream GCAUG is given in inset with colors corresponding to the key (FE test, two-tailed).

To determine the extent to which binding of CELF2 correlates with splicing, we overlaid our newly-identified functional targets of CELF2 with the CELF2 binding sites we have previously defined in Jurkat cells (Ajith et al. 2016). Consistent with our prior analysis of the more limited set of CELF2-regulated exons (Ajith et al. 2016), we find that CELF2 binds around more cassette exons that it regulates (Fig. 1C), compared to exons that are not regulated by CELF2, suggesting a direct role of CELF2 in regulation of many of the CELF2-responsive exons. However, for two-thirds of CELF2-responsive exons we find no evidence for CELF2 binding in locations that are commonly associated with splicing regulation (within the regulated exon, flanking constitutive exons or 300 nts of flanking intron, Fig. 1C). While we cannot rule out that in some of these instances CELF2 functions from a distance of greater than 300 nucleotides, or binds in a manner that is not detected by crosslinking, these results strongly suggest that CELF2 regulates a sizeable number of exons by an indirect mechanism.

### RBFOX motifs and binding are enriched downstream of CELF2-repressed exons

To investigate potential indirect mechanism for CELF2 regulation of splicing, we looked for enriched pentamers within 300 nucleotides upstream or downstream of CELF2 repressed or enhanced exons using a hypergeometric test (Fig 1D, Supplemental Table S2). Consistent with previous findings where CELF2 was shown to repress exon inclusion when bound upstream of cassette exons (Han and Cooper 2005; Dembowski and Grabowski 2009; Ajith et al. 2016), we see specific enrichment of UG-rich pentamers upstream of CELF2 repressed exons (Fig. 1D, left, blue circles). Strikingly, by contrast, we found that the top two pentamers (UGCAU and GCAUG) enriched downstream of CELF2 repressed exons (Fig. 1D, right, orange circles) are perfect matches to the high-affinity binding site of the RBFOX family, [U]GCAUG (Ponthier et al. 2006). Notably, this enrichment of RBFOX pentamers was specific to the downstream intron, from where RBFOX typically functions as a potent enhancer of exon inclusion (Yeo et al. 2009). While several other pentamers are also enriched around both CELF2 enhanced and repressed exons (Supplemental Table S2), we focused further investigation on the RBFOX motif for two reasons. First, the enrichment of the RBFOX motif downstream of CELF2-repressed exons is among the most significant, even more so than that of the UG-rich CELF2 binding sites upstream of these same exons (Fig. 1D). Secondly, the CELF and RBFOX families have both been implicated in the development and maintenance of normal function of heart, muscle, and brain (Zhang et al. 2002; Kalsotra et al. 2008; Gehman et al. 2012; Singh et al. 2014), but have yet to be directly studied in connection to one another.

We next utilized splicing code models (Barash et al. 2010) through the AVISPA tool (Barash et al. 2013) to predict the relevance of the RBFOX motifs in determining splicing outcome.Consistent with the pentamer enrichment described above, AVISPA found the RBFOX motif occurred significantly more often downstream of CELF2 repressed exons (36.6%) compared to unresponsive exons (25.6%, Fisher's exact two-tailed *p* < 3×10^−4^) or CELF2 enhanced exons (24.1%, *p* < 1 × 10^−3^, Fig. 1E inset). Moreover, the presence of the RBFOX motif downstream CELF2 repressed exons is predicted by AVISPA to have significantly more impact on splicing (higher normalized feature effect (Barash et al. 2013), see Supplemental Methods) when compared to either unresponsive (Kolmogorov-Smirnov two-sample *p* < 6.3×10^−11^) or CELF2 enhanced exons (*p* < 2×10^−4^, Fig. 1E). The RBFOX motifs downstream of the CELF2 repressed exons are also more highly conserved than those around CELF2-unresponsive exons (Fig 1F), which is an additional hallmark of functional relevance (Lambert et al. 2014; Taliaferro et al. 2016).

Finally, we utilized publicly available eCLIP peaks for RBFOX2 from a variety of cell types including HEK293, human embryonic stem cells (H1ES), HepG2, and K562 cells (Conway et al. 2016; Sundararaman et al. 2016; Van Nostrand et al. 2016) to see if there was evidence of *in vivo* binding of RBFOX2 in regions proximal to CELF2 regulated cassette exons. In line with the sequence enrichment data, we find enrichment of RBFOX2 eCLIP binding evidence within 300 nucleotides downstream of the CELF2 repressed exon, compared to unresponsive exons, in all four cell types examined (from ~14% to 25%, Fig. 2A, top). Although there was significant enrichment of RBFOX2 eCLIP peaks in most regions proximal to and within CELF2 repressed exons, the most striking enrichment in all cell types was for binding downstream of the cassette exon (Fig. 2A, bottom). This enrichment of RBFOX2 binding downstream of CELF2 repressed exons is unique compared to other RBPs analyzed by eCLIP and iCLIP experiments from ENCODE (Sundararaman et al. 2016), including CELF2 (Fig. 2B). Lastly, we investigated the binding profiles of RBFOX2 around CELF2 regulated exons on a nucleotide basis, by mapping the occurrence of high-confidence RBFOX2 eCLIP peaks (see Methods) across the sets of CELF2 regulated and unresponsive exons. In this analysis, we again observe a striking enrichment of RBFOX2 binding specifically downstream of CELF2 repressed exons (Fig. 2C). Taken together, the motif enrichment, increased splicing code feature effects and conservation scores, and enrichment of RBFOX binding, suggest that RBFOX proteins play a role in the regulation of exons that are repressed in the presence of CELF2. In particular, the enrichment of RBFOX2 binding downstream of the 5'ss, a region that has been previously identified as a location from which RBFOX2 acts as a strong enhancer of exon inclusion (Yeo et al. 2009), predicts that RBFOX proteins likely enhance the inclusion of exons deemed to be CELF2-repressed.

**Figure 2:**
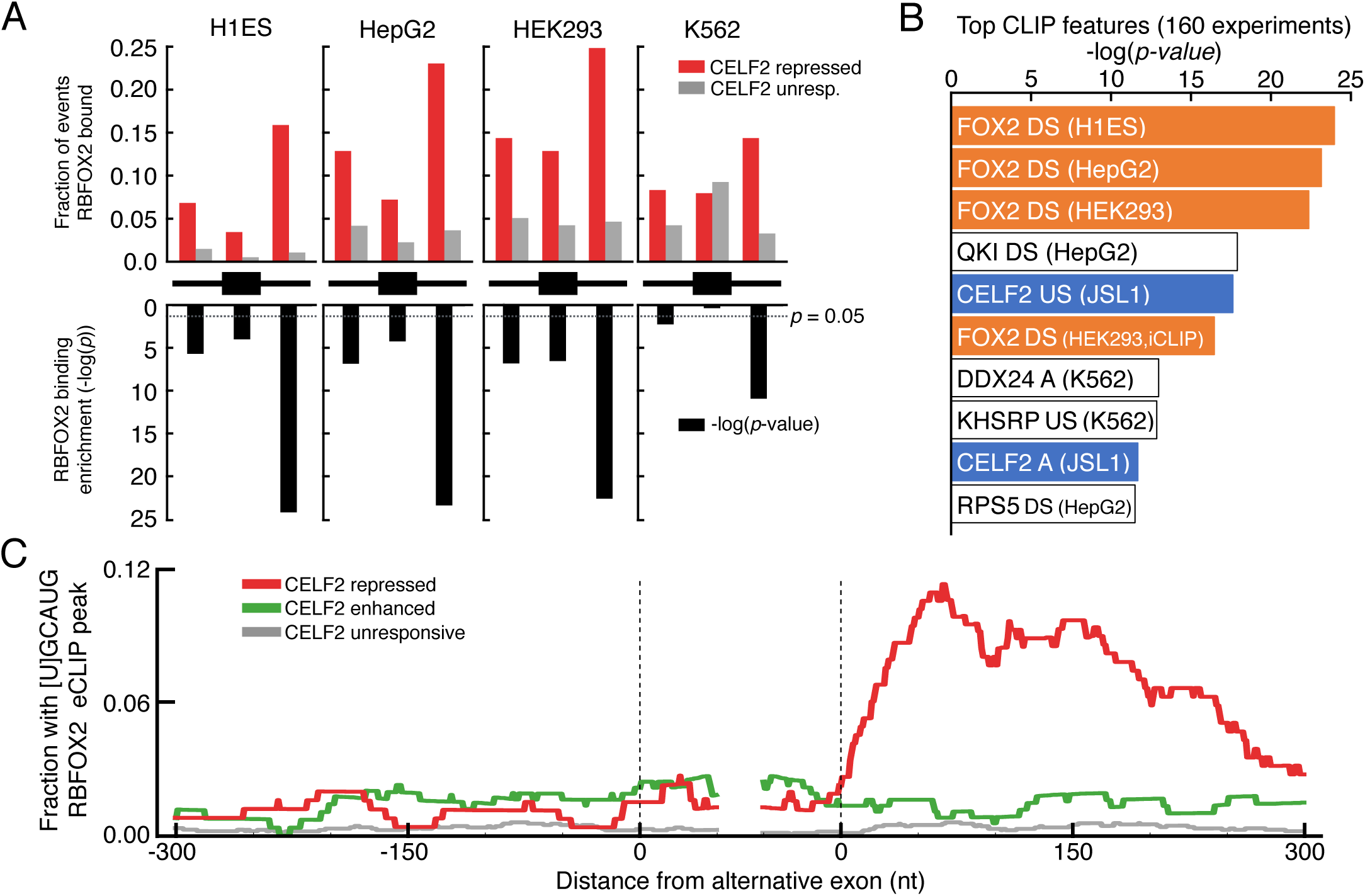
CELF2 repressed exons are highly enriched for downstream *in vivo* binding of RBFOX2. (A) Fraction of exons that were repressed by CELF2 (red) or unresponsive (grey) that contained an RBFOX2 eCLIP peak from indicated cell type in regions proximal to the alternative exon (black diagram: within 300 nt upstream, within the exon, or within 300 nt downstream) (top) and −log_10_)(*p*-value) of enrichment of RBFOX2 eCLIP peak occurrence in the CELF2 repressed versus unresponsive set (bottom, black bars, Fisher's exact two-tailed *p*). (B) Top 10 enriched CLIP features found when comparing CLIP peak occurrence from 160 experiments (ENCODE eCLIP and iCLIP; (Ajith et al. 2016)) between CELF2 repressed and unresponsive exons. Conditions in which CLIP experiment was carried out is indicated in parentheses. CLIP experiments for RBFOX2 or CELF2 are indicated in orange or blue, respectively. DS: bound downstream of alternative exon; US: bound upstream of alternative exon; A bound within alternative exon. (C) RNA maps showing the per-nucleotide frequency of [U]GCAUG containing RBFOX2 eCLIP peak occurrences within 300 nt of alternative exons that are repressed by CELF2 (red), enhanced by CELF2 (green), or were unresponsive to knockdown (grey).

### CELF2 and RBFOX2 antagonize splicing patterns in T cells

To directly test the functional relevance of RBFOX binding to CELF2-responsive exons we analyzed the relative impact of CELF2 and RBFOX depletion on the splicing of these exons. Jurkat T cells express no detectable RBFOX1 or RBFOX3 mRNA (data not shown) but do express moderate levels of RBFOX2 mRNA and protein (see below), consistent with restricted expression of RBFOX1 (muscle, heart, and brain) and RBFOX3 (brain) but more widespread expression of RBFOX2 (Conboy 2016). Therefore, we analyzed the splicing of CELF2-repressed exons in Jurkat cells depleted of CELF2, RBFOX2 or both proteins. Of the 265 high-confidence CELF2-repressed exons, 127 have evidence for RBFOX family binding in the downstream intron (eCLIP peak and/or [U]GCAUG motif). For 37 of these we also detect CELF2 binding upstream (14, Fig. 3A) downstream (5, Fig. 3C) or on both sides (18, Fig. 3B) of the regulated exon, while the remaining 90 have no CELF2 CLIP peaks in the vicinity of the CELF2-responsive exon (Fig. 3D). The very few number of CELF2-repressed exons with CELF2 binding only downstream is consistent with our earlier finding that binding of CELF2 downstream of an exon correlates mostly with exon inclusion, while binding of CELF2 upstream correlates with exon skipping (Ajith et al. 2016).

**Figure 3:**
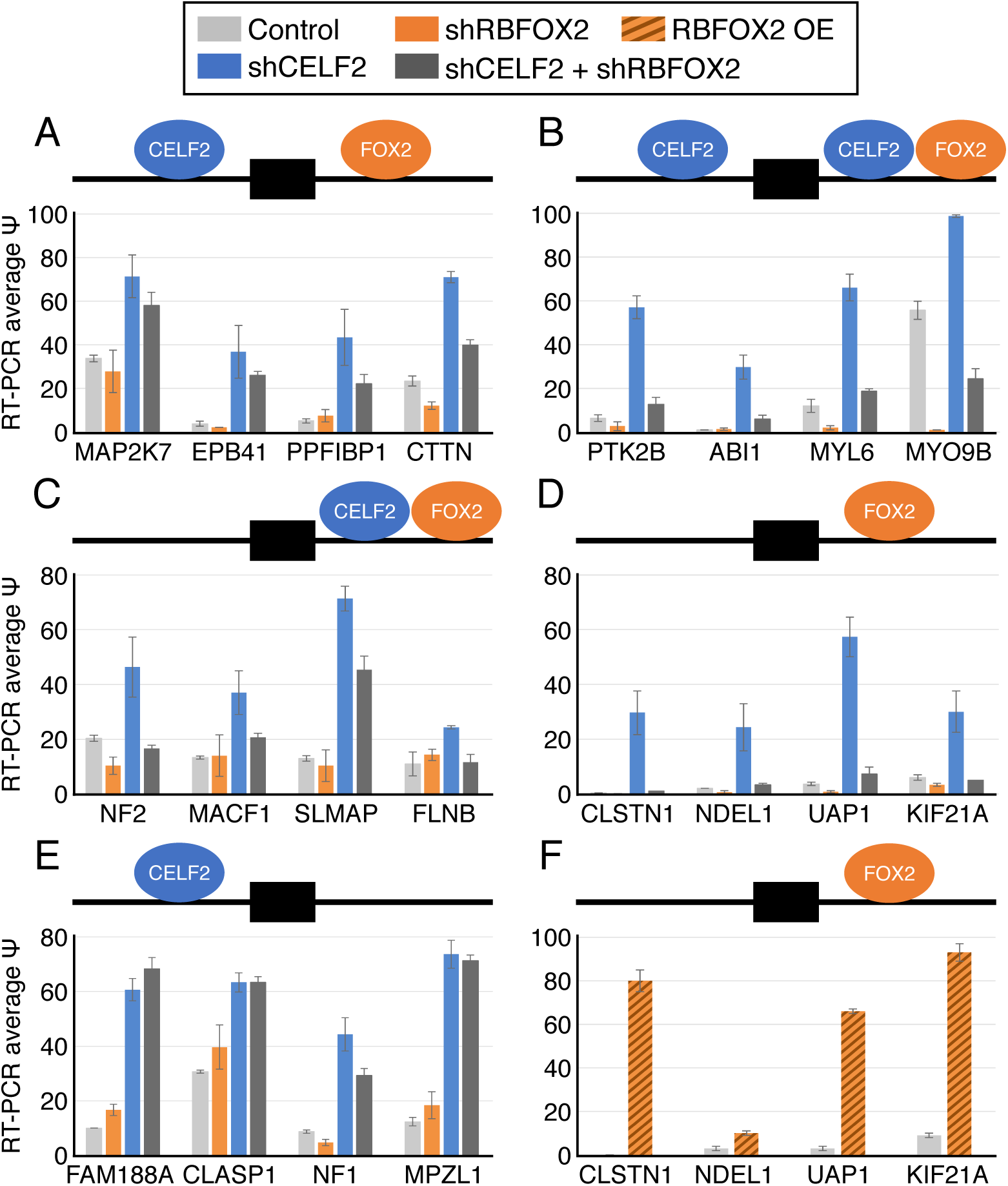
Reciprocal regulation of CELF2-repressed exons by RBFOX2 in T cells. (A-E) Quantification of splicing by RT-PCR (see Supplemental Fig. S1 for gels) of MAJIQ-identified CELF2-repressed exons in wildtype Jurkat T cells (light grey), or cells depleted of RBFOX2 (orange), CELF2 (blue) or both proteins (dark grey) by shRNA. Efficient knockdown of proteins was confirmed via Western blot (see Fig. 4). Quantifications are based on the average of no fewer than three biological replicates and error bars represent the standard deviation. Diagrams above bar charts indicate binding locations of CELF2 and/or RBFOX2 upstream or downstream of the alternative exon. Data shown is from stimulated cells, which typically express high levels of CELF2. Data for unstimulated cells is provided in Supplemental Fig. S2. (F) Quantification of splicing by RT-PCR in wildtype Jurkat T cells (light grey) or those overexpressing RBFOX2 cDNA (orange stripe). Protein expression and RT-PCR gels are in Supplemental Fig. S3.

Notably, regardless of the presence or absence of detectable CELF2 or RBFOX2 binding sites, all 20 of the 265 high-confidence CELF2-repressed exons tested show increased inclusion upon depletion of CELF2 by shRNA (Fig. 3A-E, Supplemental Figs. S1, S2), validating our identification of CELF2-repressed exons from the RNA-Seq data. Interestingly, depletion of RBFOX2 alone does have an opposite impact from CELF2 on these exons, consistent with functional antagonism; however, this impact of RBFOX2 depletion is only notable for a handful of exons compared to wildtype cells (ex. MYL6, MYO9B). By contrast, in the co-depletion of RBFOX2 and CELF2 we observe a striking antagonistic relationship that can be grouped into two patterns. For those exons with binding of CELF2 upstream, where we predict direct repression by CELF2, co-depletion of RBFOX2 has a notable but modest impact on splicing relative to CELF2 alone (Fig. 3A, blue vs dark grey), and depletion of CELF2 has a significant impact on exon inclusion even in the absence of RBFOX2 (Fig 3A, orange vs dark grey). On the other hand, for those exons around which we detect no binding of CELF, co-depletion of RBFOX2 completely abrogates the effect of CELF2 depletion (Fig 3D, light grey vs dark grey or blue), indicating that the impact of CELF2 depletion is primarily mediated indirectly via RBFOX2 (see below). Exons bound on either side by CELF2 exhibit a pattern most similar to the exons with no CELF2 binding, suggesting that perhaps in this case the enhancing and repressing activities of CELF2 cancel each other out and splicing is ultimately directed by RBFOX2 (Fig 3B). By contrast, exons that have binding sites for both CELF2 and RBFOX2 downstream show a mixed response of co-depletion, with most events resembling RBFOX2-driven splicing (Fig 3C, NF2, MACF1 and FLNB) consistent with CELF2 not typically repressing from a downstream location, while other events show a more complicated response to co-depletion (Fig 3B, SLMAP). Importantly, we observe no impact of RBFOX2 depletion on CELF2-repressed exons that lack evidence for RBFOX2 binding (Fig 3E).

In sum, we conclude that for CELF2 -bound and -repressed targets CELF2 is the primary driver of splicing, but many of these genes are also bound and enhanced by RBFOX2, which partly counters the repressive activity of CELF2. Critically, we also demonstrate that a large percentage (at least 90 of the 265 or 34%) of CELF2-repressed exons are unlikely to be direct targets of CELF2, but rather are regulated primarily through RBFOX2. Consistent with this interpretation, overexpression of RBFOX2 phenocopies knockdown of CELF2 for all 4 of the CELF2-repressed genes tested for which we have no evidence of CELF2 binding (Fig 3F vs 3D and Supplemental Fig S3).

### CELF2 represses RBFOX2 mRNA and protein levels in T cells

One mechanism that could explain the remarkable impact of RBFOX2 on indirect targets of CELF2 regulation would be if depletion of CELF2 alters the expression of RBFOX2 in Jurkat cells. Consistent with this model, analysis of *RBFOX2* mRNA expression upon CELF2 depletion by RNA-Seq revealed upregulation in *RBFOX2* mRNA levels of greater than 2.5-fold in both unstimulated and stimulated Jurkat cells upon depletion of CELF2 (adjusted *p* < 2.3×10^−11^, Fig. 4A and B). Western blotting confirmed this upregulation of RBFOX2 is also manifest at the protein level upon CELF2 knockdown (Fig. 4C). This increase in RBFOX2 protein upon CELF2 depletion occurs both in unstimulated cells, in which CELF2 expression is low, and in stimulated cells in which CELF2 expression is higher (Fig 4C, compare lanes 1 and 2 for unstimulated (left) and stimulated (right) cells). Notably, CELF2 expression is sufficient to regulate RBFOX2, independent of cell-type, as overexpression of FLAG-tagged CELF2 reduced RBFOX2 expression both in Jurkat cells as well as HEK293 cells. However, the antagonistic relationship between CELF2 and RBFOX2 expression is unidirectional, at least in Jurkat cells, as neither depletion nor overexpression of RBFOX2 caused an appreciable change in CELF2 protein (Fig. 4C, compare lanes 1 and 3 for both and Supplemental Fig. S3).

**Figure 4:**
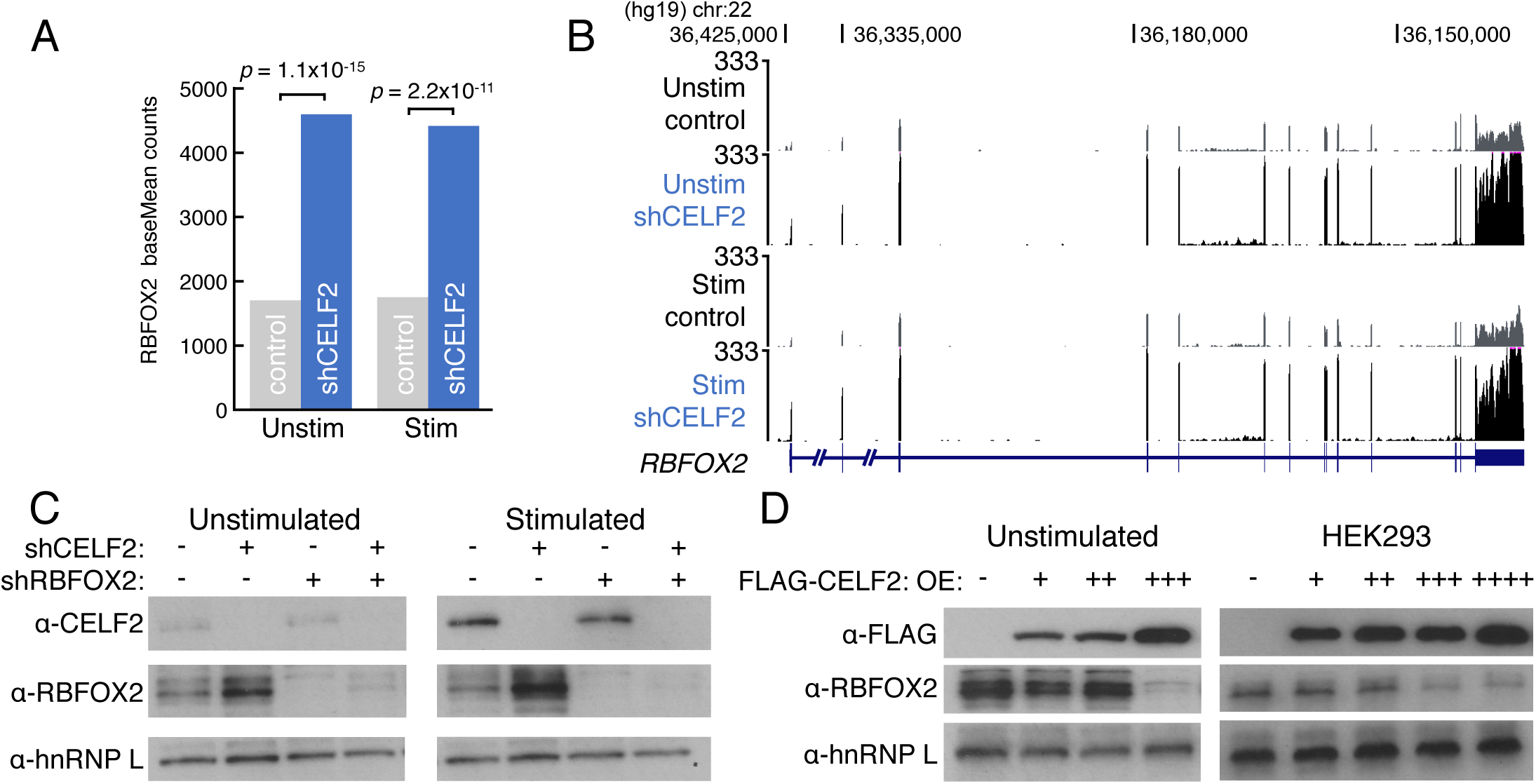
CELF2 represses RBFOX2 mRNA and protein levels in T cells. (A) RBFOX2 expression in unstimulated and stimulated JSL1 T cells (grey) compared to CELF2 knockdown cells (blue) as determined by DESeq. Significant increases in *RBFOX2* mRNA levels upon CELF2 depletion was determined by DESeq (fold-change > 2.5 and adjusted *p* < 2.3×10^−11^) (B) UCSC genome browser snapshot of RNA-seq reads over the *RBFOX2* locus in unstimulated (top) or stimulated (bottom) control (grey) versus CELF2 knockdown (blue) JSL1 cells. (C) Western blot monitoring protein levels of CELF2 and RBFOX2 in unstimulated (left) or stimulated (right) JSL1 cells depleted for CELF2 (2nd lanes), RBFOx2 (3rd lanes), or both (4th lanes). An antibody against hnRNP L was used as a loading control (D) Western blot monitoring FLAG-CELF2 and RBFOX2 expression in unstimulated JLS1 cells (left) or HEK293 cells (right) that expressed increasing amounts of FLAG-CELF2 construct. An antibody against hnRNP L was used as a loading control.

RBFOX2 is known to contain several alternative splicing events, including exon skipping of part of the single RRM which creates a dominant-negative version of the protein and an event in the C-terminal domain that can alter localization (Underwood et al. 2005; Damianov and Black 2010). Therefore, we initially reasoned that CELF2 may regulate the splicing of RBFOX2 to control expression. However, analysis of RNA-Seq data revealed no significant change in either RBFOX2 splicing event upon CELF2 knockdown (Supplemental Fig. S4A). On the other hand, we do find a marked increase in the stability of RBFOX2 mRNA in CELF2-depleted versus wildtype Jurkat cells (Supplemental Fig S4B). Thus, control of mRNA stability may provide a mechanism for CELF2-regulation of RBFOX2 expression (see Discussion).

### CELF/RBFOX antagonism is conserved in mouse heart and muscle

Given the striking functional antagonism between CELF2 and RBFOX2 in Jurkat T cells, we wished to explore if this antagonism was also evident in heart and muscle; systems in which both the CELF and RBFOX families of splicing factors are well studied and functionally critical. Previous work has shown that the CELF1 and CELF2 promotes embryonic splicing patterns in the murine muscle and heart which is reversed when both proteins are dramatically downregulated postnatally (Ladd et al. 2005; Kalsotra et al. 2008). On the other hand, RBFOX1 and RBFOX2 have been implicated as key in promoting splicing changes during myogenesis and across heart and muscle development (Gallagher et al. 2011; Gehman et al. 2012; Singh et al. 2014; Pedrotti et al. 2015), yet these two protein families have not been studied in relation to one another in these contexts.

We processed publically available RNA-seq data from adult mouse hearts that ectopically expressed CELF1 or CELF2 (Giudice et al. 2014; Wang et al. 2015) using MAJIQ and identified CELF-regulated cassette exons (see Methods). We then analyzed these cassette exons for enriched sequences and CLIP-seq features in the same way as was done in the analysis of human T cell data above (Fig. 1, 2). In line with the results from T cells, we found enriched occurrences of perfectly conserved [U]GCAUG motifs and Rbfox2 CLIP-seq peaks downstream of from 12% to 18% of CELF2 or CELF1 repressed exons (Supplemental Fig. S5). The most enriched CLIP-seq feature was for downstream binding of Rbfox2 from mouse cardiomyocytes (Fig. 5A, left). RNA maps for the per nucleotide occurrence of these Rbfox2 CLIP peaks over CELF2 repressed exons showed clear enrichment downstream in a similar region as we observed for T cells (Fig. 5B). Together these results show that exons repressed by CELF1 or CELF2 in the murine heart share features consistent with the CELF/RBFOX antagonism we observed in T cells.

**Figure 5:**
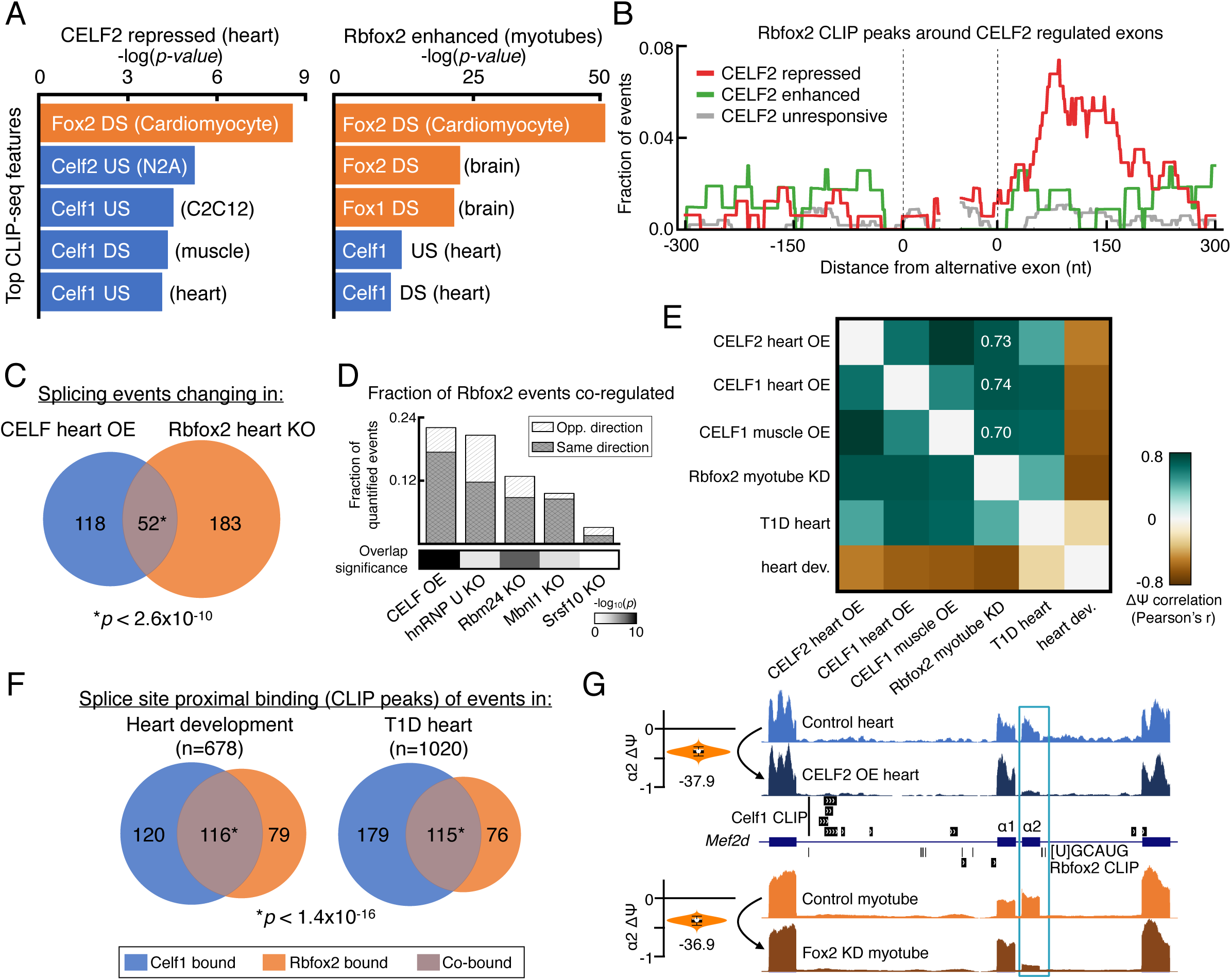
CELF/RBFOX antagonism is conserved in mouse heart/muscle and influences developmental and disease splicing programs. (A) Two-tailed Fisher's exact test −log_10_(*p*-value) for top 5 CLIP-seq peak occurrence enriched proximal to alternative exons repressed by CELF2 in mouse heart (left) or enhanced by Rbfox2 in myotubes (right) compared to unresponsive exons. Condition in which CLIP experiment was carried out is indicated in parentheses. CLIP experiments for the CELF or Rbfox families are indicated in blue or orange, respectively. DS: downstream; US: upstream. (B) RNA map of Rbfox2 CLIP-seq peak from cardiomyocytes proximal to CELF2 repressed (red), enhanced (green), or unresponsive (grey) exons in the heart. (C) Venn diagram showing overlap of splicing changes in the mouse hearts that over-expressed (OE) CELF1 or CELF2 versus events altered in a conditional heart KO of Rbfox2 (*p* < 2.6×10^−10^, Fisher's exact test). (D) Fraction of total Rbfox2 regulated events that were quantified in given condition that changed in the same direction (white) or the opposite direction (grey) of Rbfox2 regulation (top), with greyscale squares representing significance of the overlap between regulated events (bottom, (−log_10_(*p*-value), Fisher's exact test). (E) Pairwise E(ΔΨ) correlation for co-regulated (IE(ΔΨ)I > 20%) LSVs between indicated conditions. T1D: Type 1 Diabetes hearts (F) Venn diagrams showing number of cassette exons that contain a splice-site proximal CLIP-seq peak for Celf1 in heart (blue), Rbfox2 in cardiomyocytes (orange), or both (purple) and are regulated across heart development (left) or dysregulated in T1D hearts (right) (G) MAJIQ ΔΨ violin plot (left) and UCSC genome browser view with RNA-seq tracks and CLIP-seq peaks (right) around an example physiologically relevant event, the *Mef2d* α1/α2 mutually-exclusive exons, that is dysregulated in disease and antagonistically co-regulated by CELF2 (top) and Rbfox2 (bottom) (for more examples see Supplemental Table S3).

Interestingly, Rbfox2 enhanced cassette exons from myotube data showed strong enrichment of CELF1 CLIP peaks proximal to the regulated exons, suggesting that CELF/RBFOX antagonism may be a defining characteristic of not only CELF family regulated exons, but also make up a significant proportion of RBFOX family regulated exons. To further investigate this possibility, we looked for shared splicing changes identified by a targeted sequencing approach (RASL-seq,(Li et al. 2012)) from hearts of Rbfox2 knockout mice (Wei et al., 2015) and compared it to those identified by RNA-seq data from Celf1 or Celf2 overexpression in adult hearts (see Methods). Consistent with CELF/RBFOX antagonism being important for both CELF and RBFOX regulated exons in heart, there was a significant overlap in splicing changes (Fisher's exact two-tailed *p*<2.6×10^−10^), with 22% to 30% of events in either set being co-regulated (Fig. 5C). As a positive control, we repeated this analysis for other splicing factor knockouts in heart, including other UG-rich binding proteins hnRNP U and Rbm24, and found the CELF family to co-regulate the largest fraction of RBFOX2 events (Fig. 5D, top) with the most significant overlap of all RBPs tested (Fig. 5D, bottom).

Notably, a majority of these co-regulated events showed splicing changes in the same direction upon overexpression of CELF1 or CELF2 in the adult heart compared to knockout of Rbfox2, consistent with antagonistic co-regulation between these splicing factors (Fig. 5D, grey). To assess if such antagonism is true globally in the murine heart/muscle context we compared all co-regulated splicing changes quantified from RNA-seq. We again observe strong positive correlations between contexts in which CELF proteins were overexpressed compared to Rbfox2 depletion in myotubes (Pearson's r ≥ 0.7, Fig. 5E), suggesting that induced changes are not only in the same direction, but also of similar magnitude. Taken together, these results suggest CELF/RBFOX antagonism is conserved in murine muscle and heart, broadly contributes to the regulatory programs controlled by each family, and is unique among other splicing factors that are important regulators in this context.

### CELF and RBFOX co-bind regulated exons in heart development and disease

Given the evidence of conserved CELF and RBFOX family antagonism in mouse heart and muscle, we wished to examine if this relationship influenced splicing changes across development or in disease where both families, individually, have been implicated. In line with previous results (Kalsotra et al. 2008; Gallagher et al. 2011), both CELF family overexpression in adult tissues or Rbfox2 depletion largely reversed changes that occurred from embryonic to adult heart development (Fig. 5E). Rbfox2 has been shown to be dysregulated in the hearts of diabetic patients and a mouse model of Type I Diabetes (T1D) where it forms a dominant negative version of the protein, leading to a number of splicing changes (Nutter et al. 2016). Celf1 has also been shown to be upregulated in diabetic hearts via PKC signaling (Kuyumcu-Martinez et al. 2007; Verma et al. 2013), but the global consequences of this has not been examined. By analyzing public data we observe not only the previous finding that Rbfox2 depletion largely mirrors changes induced in a mouse model of T1D (r = 0.44, Fig 5E), but importantly demonstrate that overexpression of CELF1 or CELF2 also mirrors T1D heart dysregulation, in all cases more strongly than Rbfox2 depletion (0.45 ≤ r ≤ 0.72, Fig. 5E). Further consistent with a functional role for CELF and FOX family members, a large fraction of cassette exons that are regulated across development (n=678) or dysregulated in T1D hearts (n=1020) have evidence for splice site-proximal binding in the heart by Celf1 (34.8% of developmental, 28.8% of T1D) or Rbfox2 (28.8% of developmental, 18.7% of T1D) (Fig. 5F). Importantly, there is a significant amount of overlap between the events in both contexts (*p* < 1.4×10^−16^) such that, in most cases, a majority of the events bound by one RBP were co-bound by the other (39.1% to 60.2%) (Fig. 5F). In total, 17.1% of all the cassette exons regulated across heart development and 11.3% of those dysregulated in T1D hearts are co-bound by Celf1 and Rbfox2 (Fig. 5F). Notably, many of the genes that exhibit reciprocal regulation by the CELF and RBFOX families have previously been implicated in myogenesis and heart failure, such as *Mef2d* (Singh et al. 2014)(Fig. 5G, Supplemental Table S3). Together, these data suggest that antagonism between the CELF and RBFOX families contributes to splicing regulation in both normal heart development, where CELF proteins are downregulated, as well as in disease.

### RBFOX motifs are enriched around exons repressed by ancient CELF homologs

Beyond the regulation of splicing in mammals, both the CELF and RBFOX families are deeply conserved through evolution where they also regulate many aspects of RNA processing (Dasgupta and Ladd 2011; Conboy 2016). Specifically, the ancient homologs of mammalian CELF and RBFOX have been shown to bind the same UG-rich and [U]GCAUG motifs, respectively, as their mammalian counterparts (Dasgupta and Ladd 2011; Conboy 2016) and influence splicing in many non-mammals, including chicken, fruit flies, and nematodes (Ohno et al. 2012; Kuroyanagi et al. 2013; Blech-Hermoni et al. 2016; Conboy 2016). Therefore, we wished to examine if the splicing antagonism we observed in human and mouse was conserved more broadly across species. To this end, we used available RNA-seq data to identify cassette exons regulated by ancient CELF family homologs and searched for the [U]GCAUG motif. Exons that were repressed by ancient CELF homologs in primary embryonic chicken cardiomyocytes (CELF1, Fig. 6A) and *Drosophila* indirect flight muscles (Bruno or Arrest, Fig. 6B) showed a striking enrichment of the RBFOX family [U]GCAUG motif downstream of regulated exons. These results suggest that the antagonism between the CELF and RBFOX families we uncovered in human T cells (Fig. 1–4) and mouse muscle/heart (Fig. 5) displays ancient conservation through evolution.

**Figure 6:**
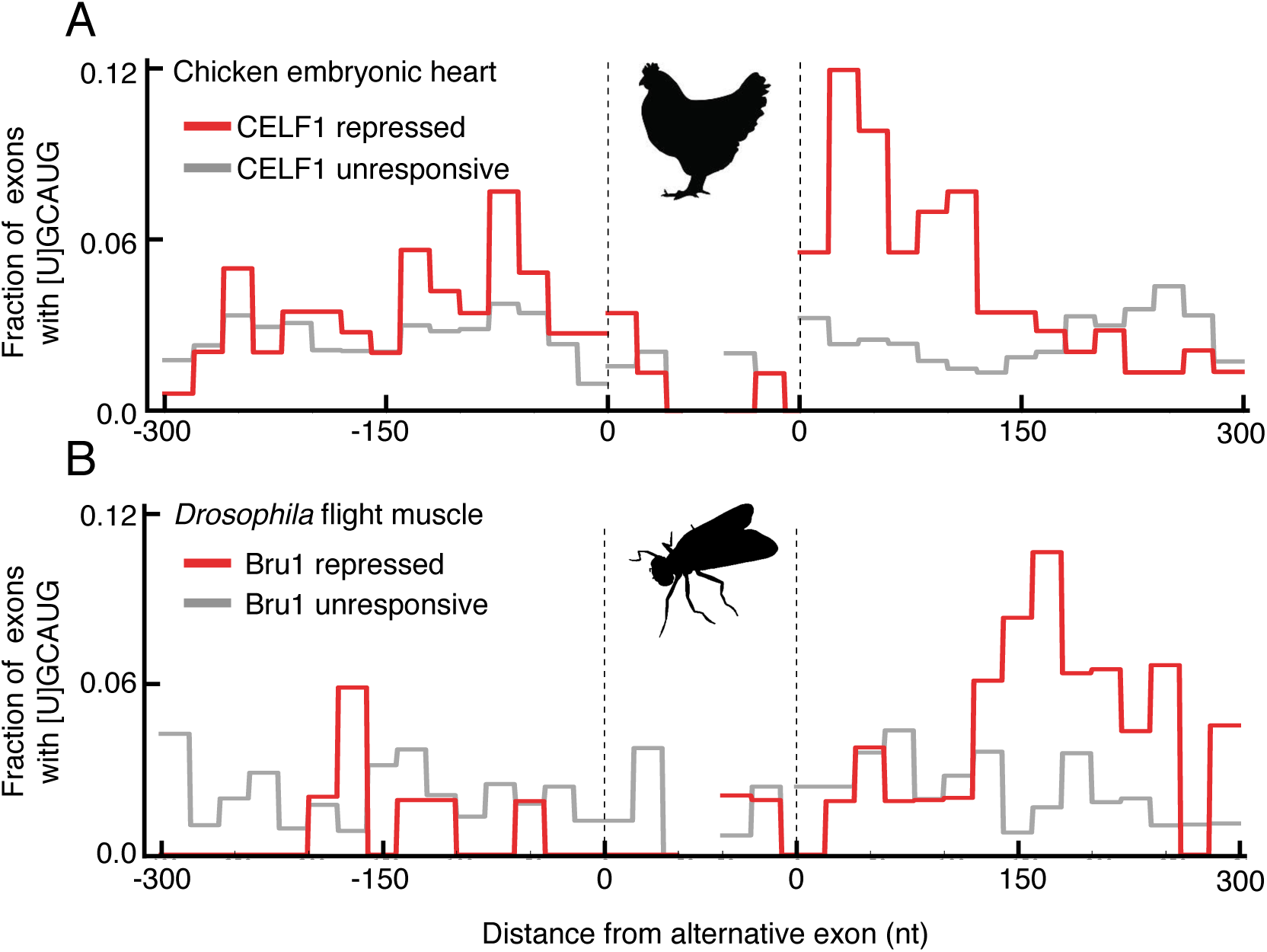
RBFOX motifs are enriched downstream of exons repressed by ancient CELF homologs. RNA maps showing the per-nucleotide (nt) frequency of [U]GCAUG motifs within 20 nt non-overlapping windows proximal to alternative exons that are repressed by ancient CELF family homologs (red) or were unresponsive to depletion/mutation (grey) in (A) chicken embryonic cardiomyocytes and (B) *Drosophila* indirect flight muscles.

## DISCUSSION

CELF2 and RBFOX2 are widely studied RNA binding proteins that both play critical roles in the regulation of tissue- and developmentally regulated splicing. In particular, both proteins have been studied extensively in neuromuscular tissues, however, cross-talk between these proteins has not been directly investigated. In studying the functional impact of CELF2 expression in Jurkat T cells, we find evidence for widespread antagonism of CELF2 and RBFOX2 in regulating splicing (Fig. 1 and 2). Some of this antagonism is through binding of both CELF2 and RBFOX2 to individual target substrates, while for other substrates the reciprocal impact of CELF2 and RBFOX2 can be primarily attributed to the fact that CELF2 negatively regulates the expression of RBFOX2 (Fig. 3 and 4). Importantly, we show that regulation of RBFOX2 expression by CELF2 is not cell-type specific, but rather also occurs in at least HEK293 fibroblasts. By analyzing publically available data, we also find evidence for antagonism between CELF2 and RBFOX2 in heart and skeletal muscle in mice, as well as in Drosophila and chicken (Fig. 5 and 6). Therefore, we conclude that antagonism between CELF2 and RBFOX2 is a conserved RBP network feature that is highly conserved throughout evolution.

### CELF2 represses expression of RBFOX2, while RBFOX2 counters the function of CELF2

Antagonism between two RBPs can occur at either the level of function or expression. Functional antagonism is seen when two RBPs both bind to the same substrate and exert opposing splicing outcomes (i.e. exon inclusion vs skipping). Such functional antagonism often involves competitive binding in which binding of one RBP prevents the binding and thus function of the other, as seen in the PTB/nPTB switch in the regulation of the N1 exon of src (Markovtsov et al. 2000). Alternatively, two RBPs may bind simultaneously to a substrate and exert opposing influences on the spliceosome (Fu and Ares 2014). In either case of functional antagonism, the protein with the greater expression or affinity typically overcomes the effect of the other protein(s). Antagonism at the level of expression, either together with functional antagonism or alone, also plays an important role in tuning RBP repertoire and activity in a cell. The genes encoding most RBPs are subject to extensive alternative splicing and other RNA processing events. Given this extent of RBP-directed processing it is not surprising that autoregulation, and trans-regulation of one RBP by others, is highly common (Boutz et al. 2007; Huelga et al. 2012; Fu and Ares 2014).

The antagonism between CELF2 and RBFOX2 involves both functional and expression antagonism. Most notably, we show that CELF2 antagonizes expression of RBFOX2 protein in both Jurkat and HEK293 cells. At least one mechanism for CELF2-mediated repression of RBFOX2 appears to be through control of mRNA stability, as both the steady-state level and half-life of RBFOX2 mRNA are increased upon knockdown of CELF2. Interestingly, we recently found that the stability of CELF2 mRNA is regulated by JNK signaling in activated T cells, which in turn is augmented by CELF2 (Martinez et al. 2015). Whether the stabilization of RBFOX2 and CELF2 mRNAs is mechanistically similar remains to be investigated. In addition, we cannot fully rule out that additional layers of regulation, including transcription and/or splicing, don’t also contribute to CELF2-mediated repression of RBFOX2 expression.

Regardless of mechanism, the antagonism of RBFOX2 expression by CELF2 clearly has a broad impact on gene expression. In particular, we identified at least 90 genes that exhibit changes in splicing upon CELF2 depletion, lack any evidence for binding of CELF2 in the vicinity of the regulated exon, and have clear evidence of RBFOX2 binding. We conclude that these 90 genes are primarily regulated by RBFOX2 rather than CELF2, as co-depletion of RBFOX2 with CELF2 fully abrogates the impact of CELF2 knockdown and direct overexpression of RBFOX2 (with no alteration in CELF2 expression) phenocopies the splicing effect observed upon knockdown of CELF2. In addition to these 90 genes regulated by expression antagonism between CELF2 and RBFOX2, we find another ~40 genes that are likely also regulated by functional antagonism of these two RBPs. These ~40 genes have evidence for binding by both CELF2 and RBFOX2, and exhibit distinct levels of exon inclusion in the doubly-deficient cells versus those depleted for either CELF2 or RBFOX2 alone. For all of these genes, loss of CELF2 in the absence of RBFOX2 leads to increased exon inclusion, while loss of RBFOX2 in the absence of CELF2 results in reduced exon inclusion. Therefore, CELF2 and RBFOX2 have opposing functions when bound to these substrates, consistent with functional antagonism. Interestingly, antagonism of CELF2 by RBFOX2 appears to be limited to this functional antagonism, as we find no evidence that RBFOX2 impacts the expression level of CELF2.

In addition to regulating splicing of the genes identified in this study, antagonism between RBFOX2 by CELF2 may play a larger role in shaping RBP expression and tissue specific splicing patterns in a range of cell types. For example, antagonism of expression may help explain the bias in CELF2 vs RBFOX2 expression that has been observed in several tissues. At one extreme, CELF2 is highly expressed in thymus/immune tissues where the RBFOX proteins show minimal expression ((Mallory et al. 2011; Mallory et al. 2015) and this study), while at the other end of the spectrum heart and skeletal muscle downregulate CELF protein expression during development but express abundant levels of RBFOX proteins (Kalsotra et al. 2008; Singh et al. 2014). Notably, both CELF2 and RBFOX2 are highly expressed and nuclear localized in the brain (Otsuka et al. 2009; Lee et al. 2016). While it is unclear how RBFOX2 avoids downregulation by CELF2 in neurons, functional antagonism between CELF2 and RBFOX2 may help explain the subset of RBFOX targets that show differential splicing between muscle and brain (Zhang et al. 2008). A number of events examined here (MAP2K7, MEF2D, EPB41) are entirely consistent with this model in that they show high levels of exon inclusion in heart, but lower in brain (MG and YB, unpublished).

### Evidence for broader competition between CELF and RBFOX families

In this study we have focused on the interplay of CELF2 and RBFOX2 with regards to alternative splicing, as these are the most highly expressed members of the CELF and RBFOX families in Jurkat cells. However, we predict the antagonism we observe here extends to other CELF and RBFOX family members. Indeed, in Figure 5 we show that splicing changes induced by overexpression of CELF1 in muscle and heart correlate as well as CELF2 overexpression with RBFOX2 knockdown. Moreover, antagonism between the CELF and RBFOX families likely extends to mechanisms other than alternative splicing. For example, recent work from the Black and Martin groups have revealed a role for cytoplasmic RBFOX1 in the brain in regulating transcript stability and/or translation (Lee et al. 2016). Notably, CELF4 is also highly expressed and localized to the cytoplasm in neurons (Wagnon et al. 2012). Analysis of iCLIP from mouse brain for CELF4 (Wagnon et al. 2012) shows nearly 90% of the cytoplasmic RBFOX1 binding sites within 3’UTRs (Lee et al. 2016) are also bound by CELF4 (MG and YB, unpublished observations). More generally, CELF proteins have been shown to recruit deadenylases to repress translation and destabilize mRNAs; whereas RBFOX promotes polyadenylation, stability and translation (Wagnon et al. 2012; Lee et al. 2016). Therefore, interplay of CELF and RBFOX proteins may have a broad role in shaping gene expression at multiple stages of RNA processing throughout many tissues and developmental conditions.

### Cooperation and Competition with other RBPs is a common feature of CELF and RBFOX families

Antagonism between CELF2 and RBFOX2 is also not the only example of functional interplay between these proteins and other RBPs. From the earliest studies linking CUG repeat expansion to myotonic dystrophy, much work has focused on antagonism between the CELF and Muscleblind (MBNL) proteins (Pascual et al. 2006). CELF and MBNL proteins primarily exhibit functional antagonism in competing for binding to CUG/UG repeats, although evidence for antagonistic expression exists as well (Wang et al. 2015). Interestingly, more recent work has shown cooperativity between MBNL and RBFOX proteins in iPSC reprogramming and muscle differentiation as well as in a myotonic dystrophy model (Venables et al. 2013; Klinck et al. 2014). RBFOX proteins have also been shown to regulate splicing in cooperation with other individual RBPs such as ESRP (Dittmar et al. 2012), PTBP (Li et al. 2015), hnRNP H (Sun et al.; Mauger et al. 2008), as well as through the activity of a large multi-protein assembly (LASR, (Damianov et al. 2016)). In sum, the antagonistic relationship between CELF2 and RBFOX2 that we describe here is only one of many positive and negative interactions between these and other RBPs that, together, shape the ultimate gene regulatory landscape of cells.

### Implications of CELF2/RBFOX2 antagonism for understanding human disease

Finally, mis-regulation or mis-expression of both CELF2 and RBFOX2 have been implicated as contributors to human disease including cancer, muscular dystrophies, heart failure and type 1 diabetes (Kuyumcu-Martinez et al. 2007; Cooper et al. 2009; Verma et al. 2013; Nutter et al. 2016). Our finding of mutual antagonism between these proteins suggests that disease-relevant splicing attributed previously to CELF2 or RBFOX2 may also be impacted by altered expression of the other. For example, Mef2d exon α2 is specifically included in heart and muscle and is known to be regulated by the RBFOX2 family (Singh et al. 2014). Inclusion of the α2 exon alters the phosphorylation and interaction partners of MEF2D to activate transcription of key myogenic genes during myogenesis and is dysregulated in heart failure where RBFOX family members are downregulated (Singh et al. 2014; Gao et al. 2016). Here we show that splicing of Mef2d exon α2 is also dysregulated upon overexpression of CELF2 in heart (Fig 5G). Other examples of disease relevant targets of both CELF2 and RBFOX2 are summarized in Supplementary Table S3 and include genes involved in calcium signaling and other members of the MEF2 family of transcription factors. In all, these results suggest that antagonism between these two families of splicing factors contributes to the altered splicing patterns in physiologically relevant targets in developmental or disease contexts in which members of one, or both, families are altered. More broadly our results emphasize the need to consider the interplay between RBPs when predicting or interpreting how dysregulation of one RBP may contribute to human disease and in designing potential therapeutic strategies.

## METHODS

### Cell culture

The JSL1 clonal population of Jurkat T cells (Lynch and Weiss 2000) was cultured in RPMI supplemented with 5% heat-inactivated fetal bovine serum (FBS) at 37°C in 5% CO_2_. For stimulation, the same growth medium was supplemented with the phorbol ester PMA (Sigma-Aldrich) for 48 hours (at a concentration of 20 ng/mL). HEK293 cells were cultured in DMEM with 10% heat-inactivated FBS at 37°C in 5% CO_2_. Depletion was achieved via stable lentiviral integration of a doxycycline-inducible shRNAs that targets CELF2 and/or RBFOX2. Targeted sequences are GCAGAGTAAAGGTTGTTGTTT (CELF2) and CCTTTAAATTTCTGCCTTTAAT (RBFOX2). Induction of the shRNA was achieved by treatment with 1 mg/ml doxycycline for 48 hrs. Expression of Flag-tagged versions of CELF2 or RBFOX2 was done through stable integration of doxycycline-inducible cDNA expression constructs into JSL1 or HEK293 cells with neomycin as a selectable marker, as described previously (Rothrock et al. 2003). Treatment with doxycycline was done the same as above.

### Western blots

Western blots were carried out as described previously, loading 10 μg of total protein lysates into 10% 37.5:1 bis-acrylamide SDS-PAGE gels (Melton et al. 2007). Antibodies used for Western blots were as follows: CELF2 (University of Florida Hybridoma Lab HL1889), RBFOX2, (Bethyl Labs A300-864A), hnRNP L (Abcam ab6106), and FLAG (Cell Signaling #2368).

### RT-PCR splicing validation

Total RNA was isolated using RNABee (Tel-Test, Inc.) by following the manufacturer's protocol. Low cycle RT-PCR was performed as described in detail previously (Rothrock et al. 2003) using 32P-labeled, sequence-specific primers to flanking constitutive exons. All primers used and expected product sizes are detailed in Supplementary Table S4. Gels were quantified by densitometry through the use of a Typhoon Phosphorimager (Amersham Biosciences).

### Library preparation and sequencing

We performed RNA-seq on Jurkat T cell line (JSL1) that were unstimulated or stimulated as well as cells in both of these conditions that were depleted for CELF2 through stable lentiviral transfection of shRNAs. All samples were sequenced in duplicate. The efficacy of knockdown was confirmed by Western blot. Total RNA was isolated as above and further purified by DNAse treatment and RNeasy Kit (Qiagen). Purity was confirmed by bioanalyzer (RIN >8). PolyA-selection, generation of stranded Illumina RNA-Seq libraries and sequencing (151 nt paired-end) were done by the Genomics Division of the Iowa Institute of Human Genetics (https://www.medicine.uiowa.edu/humangenetics/dna/). These reads have been submitted to NCBI Gene Expression Omnibus (GEO) and an accession number is pending.

### Mapping of RNA-Seq reads

For all datasets we mapped reads using STAR with genomes generated using a custom splicing junction database for each organism. We mapped reads to the following genome assemblies: human to hg19, mouse to mm10, chicken to galGal4, *Drosophila* to dm4, and *C. elegans* to ce10. For junction spanning reads we required at least 8 positions to map across the splice junction on both sides by using the option --alignSJoverhangMin 8. Only uniquely mapped reads were utilized for downstream analysis. All accession numbers and description of samples utilized in this study are summarized in Supplementary Table S5.

### Differential splicing and gene expression analysis

To quantify significant changes in gene expression from RNA-seq data, we utilized DESeq (Anders et al 2010) with the default parameters. Genes with an FDR < 0.01 and absolute log_2_ fold-change > 1 were considered to be differentially expressed.

To quantify splicing changes in all datasets we utilized MAJIQ and VOILA to detect local splice variations (LSVs), as we have described previously (Vaquero-Garcia et al. 2016). Significant splicing changes in this study were defined as those with at least one junction that has a change in expected inclusion (E(dPSI)) of greater than 20% in either direction. High-confidence changes were those LSVs that had at least one junction with a probability of |dPSI| ≥ 20% > 95% (Vaquero-Garcia et al. 2016). We classified cassette exons by combining evidence from single source and single target LSVs. For further detail on this and other computational analyses see Supplemental Methods.

### Motif and splicing code analysis

For motif enrichment we first defined splicing relevant regions for all responsive and unresponsive cassette exons. There regions were defined as: the upstream constitutive exon (C1); the alternative exon (A); the downstream constitutive exon (C2); and all intron sequence between C1 and A or A and C2 and within 300 nt of an exon (Barash et al. 2010; Barash et al. 2013). Counts for all 1024 possible pentamers were obtained for each of these regions and a hypergeometric test was applied to obtain significant differences in the presence or absence of a pentamer in a specific region in a regulated versus unresponsive set of exons. For the splicing code analysis we used AVISPA (Barash et al. 2013) as we have done previously to identify known and novel regulators of alternative splicing (Gazzara et al. 2014; Sotillo et al. 2015). For conservation analysis, we used conservation scores extracted by AVISPA that uses the phastCons46way, placental mammal track (human) or phastCons60way (mouse). The conservation of RBFOX motifs was calculated as the average conservation score of the core GCAUG motif. Perfectly conserved RBFOX motifs were defined as those with scores of 1.0 for all nucleotides.

### CLIP-seq analysis

For eCLIP and iCLIP data we downloaded all hg19 bed narrowPeak files from the ENCODE Portal (www.encodeproject.org) that were available as of October 2016 (Sundararaman et al. 2016). We processed additional publically available CLIP-seq datasets in human and mouse (summarized with accession numbers in Supplemental Table S6) using the same pipeline described for eCLIP data processing (Van Nostrand et al. 2016) that is used by ENCODE. Peaks called on replicate experiments were merged for downstream analysis. We defined high confidence RBFOX2 peaks as those that contained the [U]GCAUG consensus motif within the boundary of the peak coordinates in any of the four cell types.

To determine enriched CLIP features, a two-tailed Fisher's exact test was used to call significant differences between peak occurrence in regulated versus unresponsive sets of exons. For RNA maps, the fraction of events in each set of regulated or unresponsive exons with a CLIP peak was plotted for each position within 300 nucleotides of the cassette exons and the first and last 50 nucleotides of the cassette exon itself.

## ACKNOWLEDGEMENTS

We wish to thank all the members of the Barash and Lynch laboratories for helpful discussions and advice, particularly Caleb Radens and Jorge Vaquero-Garcia. We also acknowledge the Genomics Division of the Iowa Institute of Human Genetics for their assistance with the RNA-Seq. This work was funded by NIH grants R01 AG046544 (YB) and R35 GM118048 (KWL).

